# Sexually Dimorphic Role for Insular Perineuronal Nets in Aversion-Resistant Ethanol Consumption

**DOI:** 10.1101/2023.01.27.525899

**Authors:** Luana Martins de Carvalho, Hu Chen, Mason Sutter, Amy W. Lasek

**Author notes:** Correspondence, Amy Lasek. Department of Pharmacology and Toxicology, Virginia Commonwealth University, Richmond, VA 23298. **Acknowledgements:** This work was supported by the National Institute on Alcohol Abuse and Alcoholism (Grant R01 AA027231 and U01 AA020912 to AWL).

## Abstract

Compulsive alcohol drinking is a key symptom of alcohol use disorder (AUD) that is particularly resistant to treatment. An understanding of the biological factors that underly compulsive drinking will allow for the development of new therapeutic targets for AUD. One animal model of compulsive alcohol drinking involves the addition of bitter-tasting quinine to an ethanol solution and measuring the willingness of the animal to consume ethanol despite the aversive taste. Previous studies have demonstrated that this type of aversion-resistant drinking is modulated in the insular cortex of male mice by specialized condensed extracellular matrix known as perineuronal nets (PNNs), which form a lattice-like structure around parvalbumin-expressing neurons in the cortex. Several laboratories have shown that female mice exhibit higher levels of aversion-resistant ethanol intake but the role of PNNs in females in this behavior has not been examined. Here we compared PNNs in the insula of male and female mice and determined if disrupting PNNs in female mice would alter aversion-resistant ethanol intake. PNNs were visualized in the insula by fluorescent labeling with *Wisteria floribunda* agglutinin (WFA) and disrupted in the insula by microinjecting chondroitinase ABC, an enzyme that digests the chondroitin sulfate glycosaminoglycan component of PNNs. Mice were tested for aversion-resistant ethanol consumption by the addition of sequentially increasing concentrations of quinine to the ethanol in a two-bottle choice drinking in the dark procedure. PNN staining intensity was higher in the insula of female compared to male mice, suggesting that PNNs in females might contribute to elevated aversion-resistant drinking. However, disruption of PNNs had limited effect on aversion-resistant drinking in females. In addition, activation of the insula during aversion-resistant drinking, as measured by c-fos immunohistochemistry, was lower in female mice than in males. Taken together, these results suggest that neural mechanisms underlying aversion-resistant ethanol consumption differ in males and females.

## Introduction

Alcohol use disorder (AUD) is defined by the inability to stop or limit alcohol use despite adverse effects on one’s health, relationships, and occupation. Although more men than women have historically been diagnosed with AUD, the prevalence of this disorder in women has increased in recent decades. From 2002 to 2012, AUD diagnosis in women increased by 83.7% (1). Socioeconomic factors can account for some of this increase, but biological factors also play a role in the initiation of alcohol use and progression to AUD (2). Clinical and preclinical studies have demonstrated sex differences during different stages of the addiction cycle (3) and suggest that hormonal and genetic factors are involved in sex differences in addiction (4). It is necessary to understand the biological factors that contribute to sex differences in alcohol consumption to develop new treatments that will be effective in reducing alcohol drinking in both men and women.

A key characteristic of AUD is compulsive alcohol use, an inflexible behavior that persists despite negative consequences (5). This aversion-resistant phenotype is modeled in animals by pairing alcohol delivery with a foot shock or by adding bitter-tasting quinine to the ethanol solution. Animals that are willing to consume alcohol despite the threat of punishment (pain or bitter taste) are considered to exhibit compulsive-like behavior (6). Several studies have shown that female rats and mice exhibit higher levels of aversion-resistant alcohol consumption and alcohol seeking behavior than males (7–11). In terms of pharmacological targets for reducing aversion-resistant ethanol consumption, inhibition of orexin-1 receptor (12), diacylglycerol lipase (13), hyperpolarization-active NMDA receptors (14–16), and alpha-1 noradrenergic receptors (17) can reduce consumption of quinine-adulterated ethanol. However, except for studies examining inhibition of orexin-1 receptor (18), these experiments were conducted in male animals only. Thus, there is a great need to identify whether mechanisms that drive compulsive-like drinking in male rodents are also operational in females.

One brain region involved in compulsive alcohol drinking is the insular cortex (14, 17, 19–23). The insula is a sensory processing area that regulates decision-making, emotion, motivation, and aversion. It integrates information about body states and processes this information to influence emotions and behavior (24). A functional magnetic resonance imaging (fMRI) study in human heavy drinkers demonstrated that the anterior insula was activated when individuals viewed alcohol cues associated with the threat of electric shock (21). In addition, greater connectivity between the anterior insula and nucleus accumbens was observed in heavy vs. light drinkers, which was significantly associated with measures of compulsive alcohol use (21). Similar findings have been observed in rats, demonstrating that projections from the anterior insula to either the nucleus accumbens or brainstem promote aversion-resistant ethanol intake (14, 17). We previously found that disrupting specialized extracellular matrix structures, known as perineuronal nets (PNNs), that surround parvalbumin-expressing interneurons in the insula can render male mice more sensitive to ethanol adulterated with quinine (20), suggesting a role for insular PNNs in maintaining aversion-resistant drinking. It remains to be determined whether manipulating PNNs in female mice has the same effect as in males.

Sex differences in the density and intensity of PNNs have been described in the mammalian basolateral and medial amygdala (25, 26), with male rodents having increased numbers and/or intensity of PNNs in these brain regions. Similarly, juvenile male rats had more PNNs in the CA1 region of the hippocampus compared with juvenile females (27). However, PNNs have not been compared between males and females in the insular cortex. Given that PNNs in male mice regulate aversion-resistant ethanol consumption, that there are sex differences in PNNs, and that female mice exhibit higher levels of aversion-resistant ethanol consumption than males, we hypothesized that there would be sex differences in PNN structure in the insula that could account for increased aversion-resistant ethanol consumption in female mice. Specifically, we hypothesized that there would be increased PNNs intensity in the insula of female mice and decreased activation of the insula, as measured by c-fos immunohistochemistry (IHC) in females during drinking of quinine-adulterated ethanol. Consistent with our hypothesis, we found increased PNN intensity and decreased c-fos levels during aversion-resistant drinking in the insula of female mice compared to males. However, in contrast to males, disrupting PNNs in the insula of female mice did not affect aversion-resistant ethanol consumption. Together, these results indicate that sex-specific mechanisms regulate aversion-resistant ethanol consumption in male and female mice.

## Materials and Methods

### Animals

Adult male and female C57BL6/J mice at the age of 8 weeks were purchased from the Jackson Laboratory (Bar Harbor, ME). All mice were individually housed in a temperature- and humidity-controlled room with a 12-h reversed light/dark cycle (lights off at 10 am) for 2 weeks prior to beginning experiments and tested for alcohol drinking behavior from 10-13 weeks of age. Food and water were available *ad libitum.* Mice were fed Teklad 7912 diet (Envigo, Indianapolis, IN). All procedures with mice were conducted according to the National Institutes of Health *Guide for the Care and Use of Laboratory Animals* and approved by the UIC Animal Care and Use Committee.

### Ethanol drinking procedures

Mice were tested for consumption of quinine-adulterated ethanol in a modified drinking in the dark (DID) test. Mice received a two-bottle choice between water and 15% ethanol, 3 h into the dark cycle, for 4 h per day over the course of 5 consecutive days. Water and ethanol solutions were provided to mice in 10 ml clear polystyrene serological pipets truncated at the end to accommodate connection to a 2.5-inch stainless steel ball-bearing sipper tube (Ancare Corp, Bellmore, NY). On the first day, mice had access to ethanol without quinine. Over the next 4 days, quinine was added in increasing concentrations each day (50, 100, 250, and 500 μM, respectively) to the ethanol solution. Tube placements were alternated daily to avoid the confound of preference for a particular side. Ethanol consumption was calculated as g ethanol per kg of body weight in 4 h, and % preference for the ethanol solution was calculated as volume of ethanol consumed divided by total fluid consumed x 100. To test consumption of a quinine solution without ethanol, a separate group of mice were presented with a two-bottle choice of water or water plus quinine in the dark cycle for 4 h per day over 5 days. On the first day, both bottles contained water, then over the next 4 consecutive days one bottle of water had quinine added in increasing concentrations daily as described above. Quinine consumption was calculated as ml quinine solution per kg body weight over 4 h and % preference was calculated as volume of quinine solution divided by total fluid consumed x 100. Finally, a one-bottle DID experiment was done with 20% ethanol or water (control) for 4 h in one day to test the effect of acute ethanol exposure on aggrecan and brevican protein levels in the insula by western blot. Male mice consumed 3.4 ± 0.29 g/kg and females consumed 6.4 ± 0.95 g/kg ethanol during the 4 h session. Insula samples were collected immediately after the drinking session.

### PNN labeling and immunohistochemistry (IHC)

Mice were euthanized during the dark cycle with a lethal dose of a commercial euthanasia solution containing pentobarbital and then transcardially perfused with ice-cold PBS followed by 4% paraformaldehyde. Brains were post-fixed in 4% paraformaldehyde overnight and cryoprotected in 30% sucrose for 48 hours. Every 4^th^ section of 50 μm serial coronal sections were collected from the area of the brain containing the insula, spanning 1-2 mm anterior to bregma. Free-floating sections were treated twice with 50% ethanol and then treated with 1% hydrogen peroxide. Sections were blocked in carbo-free blocking solution (Vector Laboratories, Burlingame, CA), and then incubated with biotinylated *Wisteria floribunda* agglutinin (WFA, Vector Laboratories, 1:1000) to label PNNs, followed by Dylight 488-conjugated streptavidin secondary antibody (Vector Laboratories). Subsequently, the sections were washed with PBS and blocked with normal donkey serum (Jackson Immunoresearch, West Grove, PA) and then incubated with mouse anti-parvalbumin (PV, #195011, Synaptic Systems, Göttingen, Germany, 1:1000), followed by Alexa Flour 594-conjugated secondary antibody (#711-585-150, Jackson Immunoresearch, 1:4000). Fluorescent sections were mounted on slides with Fluoromount-G from SouthernBiotech (Birmingham, AL). For c-Fos IHC, 40 μm serial coronal sections were treated with 1% hydrogen peroxide for 10 min and blocked with 5% goat serum plus 0.25% Triton X-100 for 1 h at room temperature, then incubated with rabbit-anti mouse c-Fos antibody (Synaptic Systems, # 226008, 1:1000) in blocking solution overnight at 4°C. Then sections were incubated with biotinylated goat anti-rabbit IgG (Vector Laboratories, #PK4001, 1:500). DAB signal was developed by VECTASTAIN® ABC-HRP Kit and DAB Substrate Kit (Vector Laboratories, # PK4001 & SK4100).

### Image acquisition and analysis

Fluorescent images were captured using Olympus BX51/IX70 fluorescent microscope (Olympus, USA) with a 10x objective. Cell intensity and counting was performed using the ImageJ macro plug-in Pipsqueak AI (Rewire Neuro, Portland, OR) (Slaker et al., 2016). Pipsqueak AI was run in “semi-automatic mode” to select ROIs to identify individual PV+ cells and PNNs. Double-labeled neurons were considered when WFA was located around the perimeter of the PV+ staining inside the cell. A Zeiss Laser Scanning Confocal Microscope (LSM) 710 META was used for the image in Figure 4G at a 40x objective and optical thickness of 2.8 μm. For imaging c-fos IHC, brightfield images were taken with a 20x objective using a Zeiss AxioScope A1 microscope and analyzed using ImageJ.

### Western blotting

Frozen tissue punches containing the insula were manually homogenized in ice cold 1X RIPA buffer (Cell Signaling Technology, Danvers, MA) containing 1X Halt protease inhibitor cocktail (Thermo Fisher Scientific). Protein concentrations were determined using the Pierce BCA Protein Assay Kit (Thermo Fisher Scientific). Equal amounts of protein (15 μg) were separated by SDS-PAGE on precast Novex 4%–12% Tris-glycine gels (Thermo Fisher Scientific) and transferred to nitrocellulose membranes. Membranes were blocked with bovine serum albumin in Tris-buffered saline (20 mM Tris, 150 mM NaCl, pH 7.4). Membranes were then incubated with anti-aggrecan (#AB1031, Millipore Sigma, 1:1000) or anti-brevican (#610894, BD Biosciences, 1:2000) and anti-β-actin (#A5441, Millipore Sigma, 1:16,000) antibodies. Secondary antibodies were donkey anti-mouse IgG DyLight 680 (#SA5-10170, Thermo Fisher Scientific, 1:20,000) and donkey anti-rabbit IgG DyLight 800 (#SA5-10044, Thermo Fisher Scientific 1:5000). Blots were imaged on an Odyssey Fc Dual-Mode Imaging system (LI-COR) and analyzed using Image Studio Lite (LI-COR).

### Stereotaxic surgery and intra-cranial injections

For injection of chondroitinase ABC (ChABC), C57BL6/J mice at the age of 8-12 weeks were anesthetized with ketamine (100 mg/kg, intraperitoneal [i.p.]) and xylazine (10 mg/kg, i.p.) and placed into a digital stereotaxic apparatus. A 5 cm incision was made in the scalp and 0.28 mm diameter holes were drilled bilaterally in the skull. ChABC (0.5 μl of a 50 U/ml solution in PBS, Millipore Sigma, St. Louis, MO, USA) or PBS was infused at a rate of 0.2 μl per min into the anterior insular cortex (1.5 mm anterior to bregma, 2.9 mm from the midline, and 2.6 mm ventral from the top of the skull) using 33-gauge stainless steel hypodermic tubing connected to a 1 μl gastight Hamilton syringe with PE20 tubing and an infusion pump. Mice were given a subcutaneous injection of meloxicam (2 mg/kg) for analgesia immediately after the completion of the surgery. Mice recovered for 3 days prior to testing ethanol consumption.

### Experimental design and statistical analysis

Data are presented as the mean ± SEM. Statistical analyses were performed using Prism software (version 9.1, GraphPad, San Diego, CA). Two-way repeated measures (RM) ANOVAs were performed for the results presented in Figure 1 (with sex as the between-subject factor and quinine concentration as the within-subject factor). Two-way ANOVA was performed for results in Figure 2E (with sex and treatment as between-subject factors). Three-way RM ANOVAs were performed for the results presented in Figure 3 with sex and treatment as between-subject factors and quinine concentration as the within-subject factor. These were followed by two-way RM ANOVAs within each sex with quinine concentration as the within-subject and treatment as the between-subject factor. Bonferroni’s multiple comparisons tests were done if there was a significant quinine concentration by treatment interaction. The number of mice in each group is indicated in the figure legend for each figure. For analysis of the fluorescently stained sections, an unpaired student’s t-test was performed with sex as the between-subject factor. For reporting PNN intensity (Figure 2B), each data point represents the fluorescent WFA intensity in a single section, obtained by calculating the average fluorescent intensity of ~100 individual cells per section. WFA intensity and cell counts were obtained from 4 mice per sex, with 4 sections (females) and 2-4 sections (males) per mouse, giving an n of 16 for females and 13 for males. For categorizing PNNs into low, medium, and high intensity (Figure 2C), quartiles were used as the cutoff based on the intensity of PNNs in female mice. Low intensity PNNs were below the 25th percentile (greater than 5.23 raw intensity units), medium intensity PNNs were between the 25th and 75th percentile (5.23-6.65 raw intensity units) and high intensity PNNs were in the 75th percentile (greater than 6.65 raw intensity units). A Chi-squared test was performed to evaluate the distribution between sexes. For the c-fos IHC in Figure 4, data was analyzed by two-way ANOVA with sex and region as between-subject factors. Each data point is the average number of c-fos counts per mouse, with 4 sections analyzed per mouse and 6 mice per sex.

**Figure 1.**
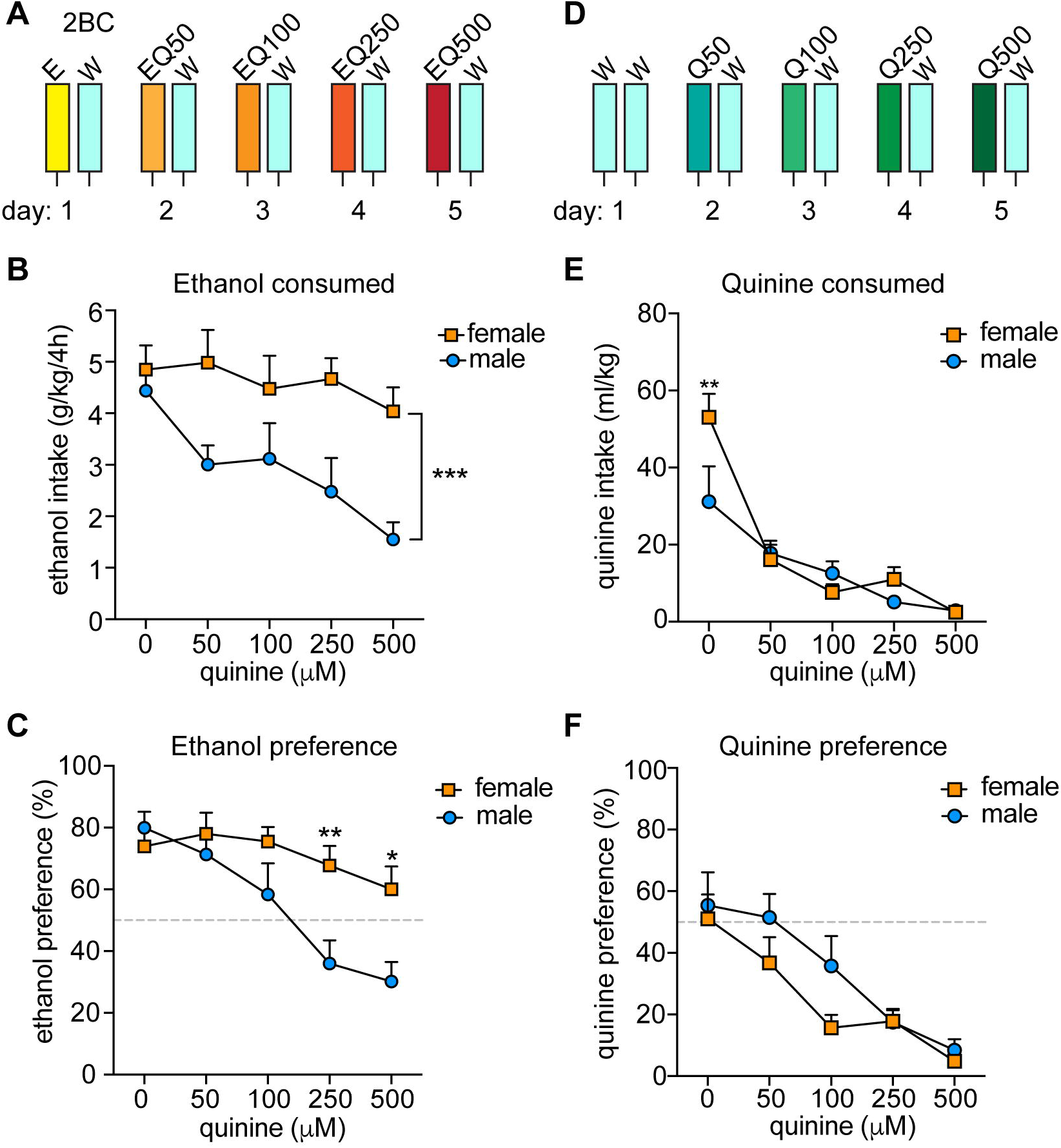
Sex differences in aversion-resistant ethanol consumption. (**A**) Experimental design for measuring aversion-resistant ethanol consumption. Mice (n = 11 males and 12 females) were given a two-bottle choice (2BC) ethanol drinking in the dark test with 15% ethanol (E) and water (W) on the first day. Each day, the quinine (Q) concentration was increased in the ethanol from 50 to 500 μM (EQ50, EQ100, EQ250, and EQ500). (**B**) Daily ethanol consumed in g per kg body weight over 4 h. ***p<0.001, effect of sex by two-way RM ANOVA. (**C**) Daily ethanol preference, expressed as ethanol consumed divided by total fluid consumed x 100. **p<0.01 and *p<0.05 when comparing females to males at EQ250 and EQ500, respectively, by Bonferroni’s post-hoc multiple comparisons testing. (**D**) Experimental setup for measuring quinine consumption. Mice (n = 8 males and 8 females) underwent 2BC in the dark on the first day with water only and then with quinine doses increasing daily from 50 to 500 μM (Q50, Q100, Q250, Q500). (**E**) Daily quinine consumed in ml quinine per kg body weight over 4 h. **p<0.01 when comparing females to males in water consumption by Bonferroni’s post-hoc multiple comparisons testing. (**F**) Daily quinine preference, expressed as quinine consumed divided by total fluid consumed x 100.

**Figure 2.**
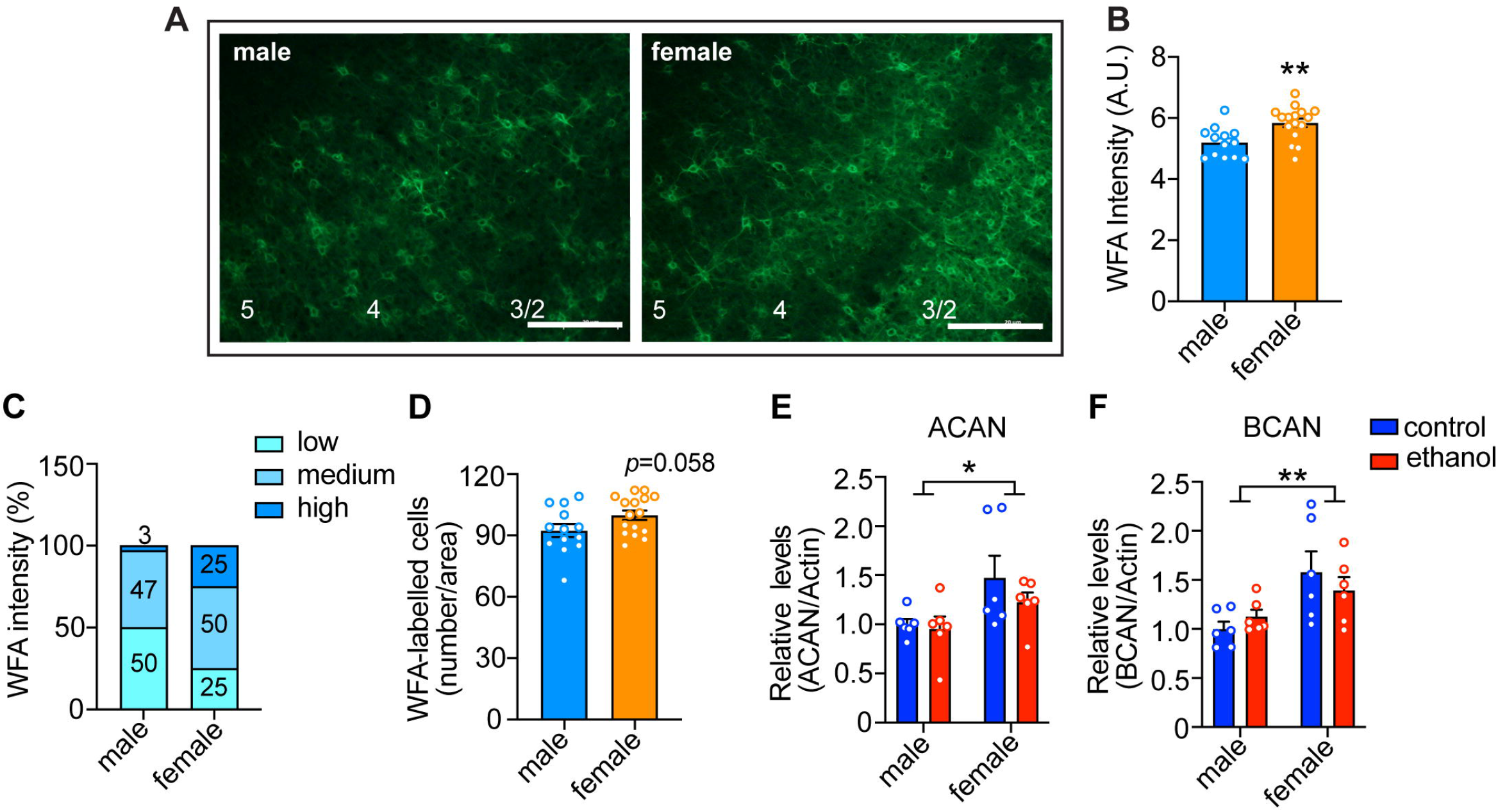
Sex differences in perineuronal net (PNN) intensity in the anterior insula. (**A**) Representative images of male (left panel) and female (right panel) insula sections that were fluorescently labelled with *Wisteria floribunda* agglutinin (WFA), which binds to PNNs. Scale bar, 200 μm. Cortical layers are indicated at the bottom. (**B)** Quantified WFA fluorescence intensity. Each data point represents the average WFA intensity surrounding neurons in a single section, with ~100 cells per section analyzed in 2-4 sections per mouse from 4 mice per sex. (**C**) Percentage of neurons with low, medium, and high intensity PNNs. There was a significant sex difference, with females having more high intensity and less low intensity PNNs than males. X^2^= 9.77 x 10^-37^. (**D**) Average number of WFA-labelled cells per section. (**E-F**) Aggrecan (**E**, ACAN) and brevican (**F**, BCAN) protein levels in male and female mice that drank ethanol (n=6 per sex) or water (n=6 per sex) for 4 h in the dark. Protein levels are shown relative to β-actin ban intensity as a loading control. *p<0.05 and **p<0.01, main effect of sex by two-way ANOVA.

**Figure 3.**
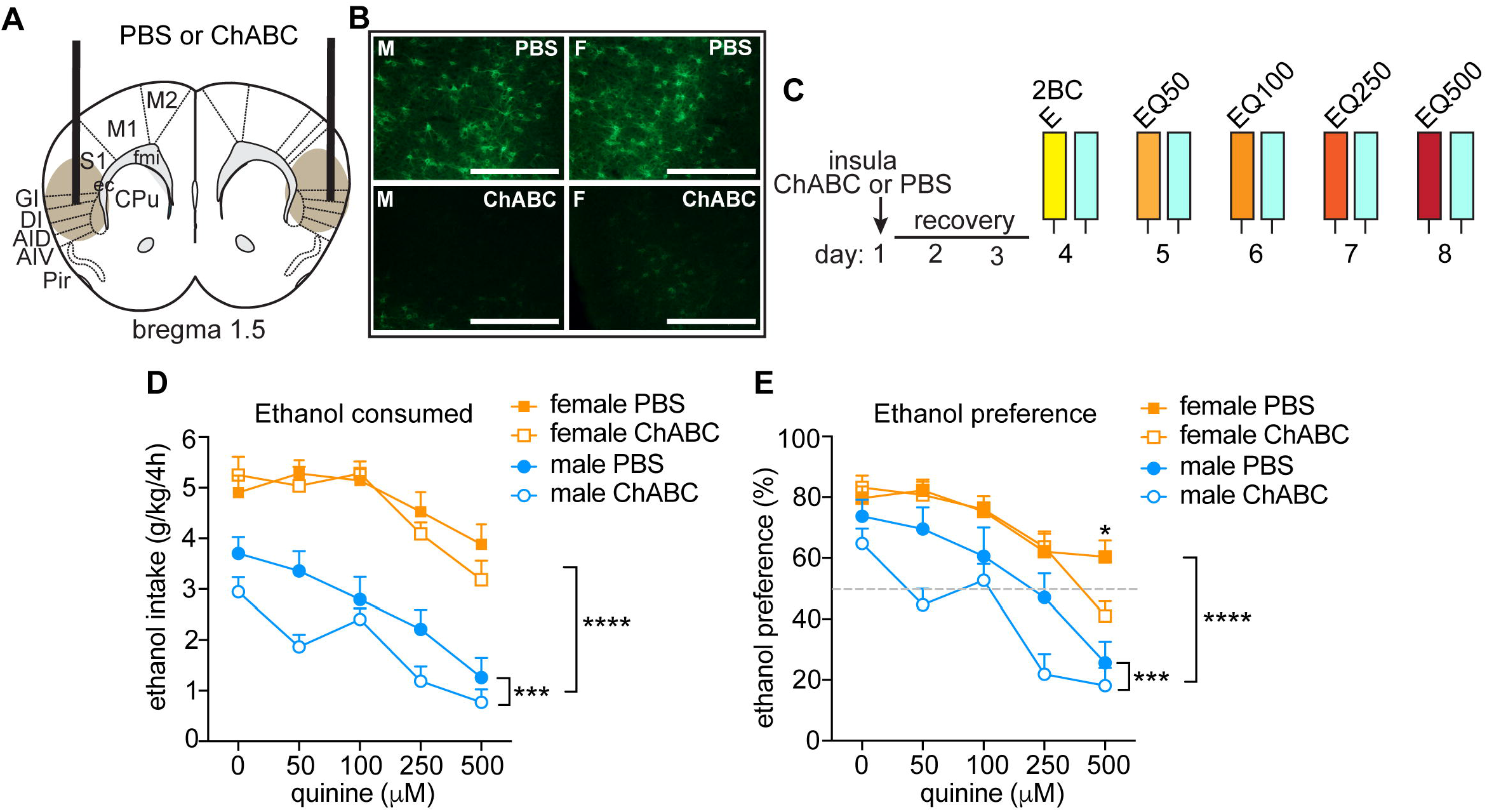
Disrupting PNNs only in male mice increases sensitivity to ethanol-adulterated quinine. (**A**) Illustration of mouse brain coronal section indicating sites of chondroitinase ABC (ChABC) or PBS injection into the anterior insula. Brown oval shows approximate spread of ChABC, as determined by reduced fluorescent WFA staining. (**B**) Representative images of fluorescent WFA staining in male (M, left panels) and female (F, right panel) mice 3 days after PBS (top panels) or ChABC injections into the insula. Scale bar, 200 μm. (**C**) Experimental design. Mice were injected with ChABC (n=16 males and 16 females) or PBS (n= 16 males and 16 females) in the insula and recovered from surgery for 3 days, followed by two-bottle choice (2BC) ethanol drinking in the dark. Quinine was added in increasing concentrations from 50-500 μM each day to the ethanol (EQ50, EQ100, EQ250, EQ500). (**D**) Daily ethanol consumed in g per kg body weight in 4 h. (**E**) Daily ethanol preference, expressed as ethanol consumed divided by total fluid consumed x 100. ****p<0.0001, main effect of sex by three-way ANOVA; ***p<0.001, main effect of ChABC in males by two-way ANOVA; *p<0.05, PBS vs. ChABC in females at 500 μM quinine.

**Figure 4.**
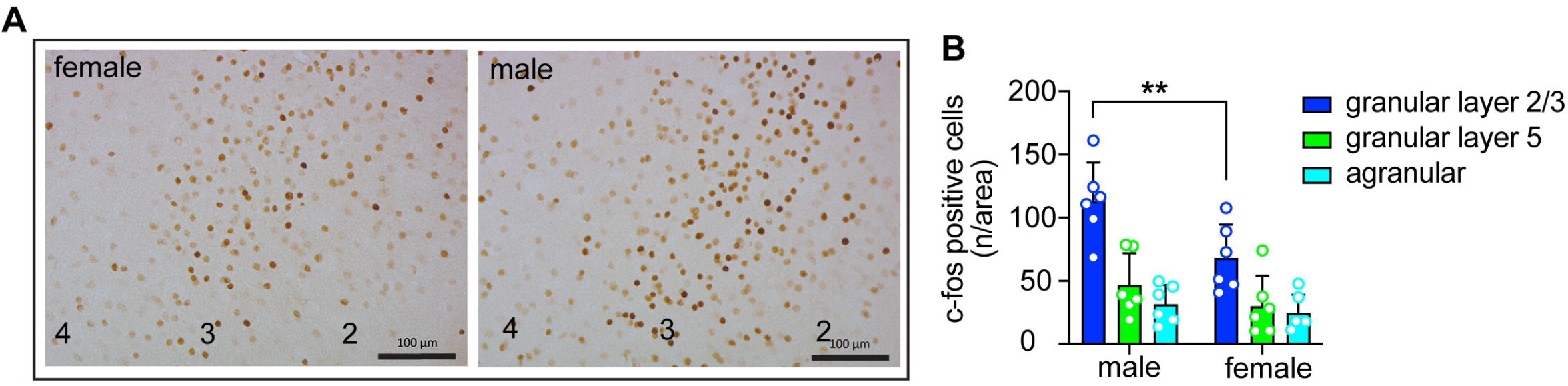
Sex differences in insula activation during consumption of quinine-adulterated ethanol. (**A**) Representative images of c-fos immunohistochemistry in the granular layer of the insula of male and female mice after drinking ethanol with 250 μM quinine. Cortical layers are indicated at the bottom. Scale bars, 100 μm. (**B**) Quantification of c-fos immunohistochemistry (n = 6 males and 6 females) in the granular and agranular layers of the insula of mice after drinking ethanol with 250 μM quinine. **p<0.05, effect of sex in granular layer 2/3 by Bonferroni’s multiple comparisons test after two-way ANOVA.

## Results

### Sex differences in aversion-resistant ethanol consumption

Previous reports indicated that female mice exhibit greater aversion-resistant ethanol consumption than males when tested for quinine-adulterated ethanol drinking (7, 10). To confirm these results, we performed a modified two-bottle choice DID procedure in which quinine was added to the ethanol solution consecutively each day in sequentially increasing concentrations (Fig. 1A; 50, 100, 250, and 500 μM quinine). We observed significant sex differences in both ethanol consumption and preference, with male mice reducing ethanol intake and preference as the quinine concentration increased, whereas females did not significantly alter their ethanol consumption or preference at any of the tested quinine concentrations (Fig. 1B, ethanol consumption: sex, *F*_(1, 21)_ = 15.16, *P* = 0.0008; quinine concentration, *F*_(4, 84)_ = 3.97, *P* =0.0054; Fig. 1C, ethanol preference: sex, *F*_(1, 21)_ = 6.66, *P* = 0.017; quinine concentration, *F*_(4, 84)_ = 11.32, *P* < 0.0001). There was also a significant interaction between sex and quinine concentration for ethanol preference (*F*_(4, 84)_ = 3.63, *P* = 0.0089; post-hoc Bonferroni’s multiple comparisons test, *P* = 0.0071 and *P* = 0.013 when comparing males and females at 250 and 500 μM quinine, respectively). These results confirm that female mice demonstrate higher levels of aversion-resistant ethanol drinking.

To determine if female mice are simply less sensitive than males to the bitter taste of quinine, we tested them in a modified two-bottle choice DID procedure with quinine added to water (Fig. 1D). Both sexes similarly decreased their consumption and preference for the quinine-adulterated water as the quinine concentration increased (Fig. 1E, consumption: quinine concentration, *F*_(4, 56)_ = 27.94, *P* < 0.0001; Fig. 1F, preference: quinine concentration, *F*_(4, 56)_ = 17.46, *P* < 0.0001). There were no significant effects of sex on water consumption or preference, although there was a sex by quinine concentration interaction for water consumption on the first day, which was due to females drinking more water relative to body weight than males (*F*_(4, 56)_ = 3.32, *P* = 0.017; post-hoc test comparing males to females at 0 μM quinine, *P* = 0.002). Thus, male and female mice appear to be equally sensitive to the taste of quinine, indicating that the increased aversion-resistant ethanol drinking by female mice is not simply due to deficient bitter taste perception.

### Sex differences in PNN intensity in the anterior insula

We previously demonstrated a role for PNNs in the insular cortex of mice in modulating aversion-resistant ethanol drinking (20). One possible reason for greater aversion-resistant ethanol consumption in females could be due to altered PNN structure in the insula. To test this, we fluorescently labeled PNNs using WFA in the insula of mice that were euthanized during the dark phase because this is when we found sex differences in aversion-resistant drinking and PNN intensity has been shown to vary throughout the diurnal cycle (28, 29). The mean WFA fluorescence intensity around individual neurons was greater in females than in males (Fig. 2A, B; *t*_27_ = 3.26, *P* = 0.003; female mean: 5.84 ± 0.14, male mean: 5.20 ± 0.13), which was primarily seen in cortical layers 2/3 (Fig. 2A). Additionally, when PNNs were categorized as high, medium, or low intensity, only 3% of the PNNs in males were high intensity compared to 25% in females, while 50% of PNNs in males were categorized as low intensity compared to 25% in females (Fig. 2C, χ^2^ = 22.71, *P* < 0.0001). No significant difference was found between males and females in the number of neurons surrounded by PNNs, although there was a trend towards an increase in females (Fig. 2D, t_27_ = 1.97, *P* = 0.058; female mean: 99.88 ± 2.29, male mean: 92.38 ± 3.12).

To confirm the sex difference in PNNs and to determine if acute ethanol exposure affects the expression of PNN proteins, we performed western blots on insula homogenates from male and female mice using antibodies to aggrecan and brevican, proteoglycan components of PNNs. Mice underwent a one-bottle DID procedure for one 4 h session with ethanol or water as a control prior to dissecting the insula. Aggrecan and brevican protein levels were significantly higher in the insula of females compared to males independently of whether mice drank ethanol or water (Fig. 2E, aggrecan: sex, *F*_(1, 20)_ = 7.03, *P* = 0.015; Fig. 2F, brevican: sex, *F*_(1, 20)_ = 9.84, *P* = 0.0052). A single ethanol drinking session did not alter aggrecan or brevican protein levels. Full images of western blots are in Supplementary Fig. 1. Together, these results indicate that females have innately higher PNN deposition around insular neurons than males during the dark phase of the diurnal cycle.

### Disrupting PNNs in male mice increases sensitivity to ethanol-adulterated quinine

To determine if disrupting PNNs in female mice alters aversion-resistant ethanol drinking, we injected ChABC, a bacterial enzyme that digests chondroitin sulfate glycosaminoglycans, into the anterior insula and compared the results to males injected intra-insula with ChABC (Fig. 3A, B). Mice were tested 3 days after injection for consumption and preference of quinine-adulterated ethanol as described above (Fig. *3*C). A three-way RM ANOVA indicated significant main effects of treatment (*F*_(1, 60)_ = 5.29, *P* = 0.025), quinine concentration (*F*_(4, 240)_ = 34.17, *P* < 0.0001) and sex (*F*_(1, 60)_ = 120.9, *P* < 0.0001) on ethanol consumption (Fig. 3D). Similar results were found for ethanol preference (Fig. 3E, treatment, *F*_(1, 60)_ = 5.74, *P* = 0.0197; quinine concentration, *F*_(4, 240)_ = 46.2, *P* <0.0001; sex, *F*_(1, 60)_ = 36.45, *P* <0.0001; treatment by quinine concentration by sex interaction, *F*_(4, 240)_ = 2.54, *P* = 0.041). Since we observed sex differences in ethanol drinking, we next performed focused two-way RM ANOVAs within each sex to determine if ChABC treatment significantly altered ethanol consumption and preference. In males, ChABC treatment resulted in reduced ethanol intake and preference when compared to the PBS control (ethanol consumption: treatment, *F*_(1, 30)_ = 5.81, *P* = 0.022; quinine concentration, *F*_(4, 120)_ = 25.66, *P* < 0.0001; ethanol preference: treatment, *F*_(1, 30)_ = 5.78, *P* = 0.023; quinine concentration, *F*_(4, 120)_ = 23.97, *P* < 0.0001), whereas in females, ChABC had no significant effect on ethanol consumption and only reduced ethanol preference at 500 μM quinine when compared to the PBS control (ethanol consumption: treatment, *F*_(1, 30)_ = 0.43, *P* = 0.52; quinine concentration, *F*_(4, 120)_ = 12.38, *P* < 0.0001; interaction, *F*_(4, 120)_ = 1.05, *P* = 0.39; ethanol preference: treatment, *F*_(1, 30)_ = 0.52, *P* = 0.48; quinine concentration, *F*_(4, 120)_ = 24.67, *P* < 0.0001; interaction, *F*_(4, 120)_ = 2.96, *P* = 0.023, post-hoc Bonferroni’s, PBS vs. ChABC at 500 μM quinine, *P* = 0.015). These results show that disrupting PNNs in the insula of male mice renders them sensitive to quinine-adulterated ethanol, while the same manipulation in the insula of female mice alters their sensitivity to quinine-adulterated ethanol only at a very high quinine concentration.

### Sex differences in insula activation during consumption of quinine-adulterated ethanol

Excitatory neurons in the insula are activated during drinking of ethanol containing quinine (30). To determine if the extent of insular neuron activation differs by sex, we measured c-fos expression by IHC in male and female mice after an ethanol plus quinine drinking session. c-fos expression was higher in granular layers 2/3 compared to granular layer 5 and agranular regions of the insula in both sexes. In addition, c-fos expression was elevated in males compared to females (Fig. 5; region, *F*_(2, 30)_ = 24.84, *P* < 0.0001; sex, *F*_(1, 30)_ = 8.69, *P* = 0.0061; post-hoc Bonferroni, *P* = 0.0064 when comparing males and females within granular layers 2/3). These results suggest greater activation of the insula of male mice compared to female mice during ethanol-adulterated quinine drinking.

## Discussion

This study confirms previous observations that female mice are more resistant than males to the suppressive effects of quinine on ethanol consumption and provides new evidence that mechanisms regulating aversion-resistant ethanol consumption differ in male and female mice. We previously demonstrated that PNNs in the insula of male mice promote aversion-resistant ethanol consumption. We show here that females have a higher intensity of PNN staining in the insula, suggesting that this might be a factor that contributes to increased aversion-resistant drinking in females. However, disrupting PNNs in females did not result in reduced intake of ethanol-adulterated quinine as it did in males.

The observation that female mice are less sensitive to the addition of quinine to ethanol than male mice are consistent with other studies. For example, Fulenwider et al found that ethanol consumption was suppressed in male mice when 30, 100, and 300 μM quinine was added to the ethanol, but 300 μM quinine was required to suppress ethanol intake by females (7). Sneddon et al showed that female mice self-administered more ethanol in an operant procedure and were able to tolerate higher quinine concentrations in the ethanol solution than males (10). However, others have reported no sex differences in quinine-adulterated ethanol intake, but testing was done after a few weeks of pre-exposure to ethanol in limited-access procedures (31, 32), which has been shown to induce aversion-resistant ethanol intake in male mice (33). Thus, innate sex differences in aversion-resistant ethanol consumption may disappear after repeated bouts of ethanol exposure but this could depend on the exposure method. We found that males and females were equally sensitive to the suppressive effect of quinine on water intake, indicating that differences in bitter taste perception are not the cause of increased aversion-resistant ethanol drinking in female mice.

We initially hypothesized that sex differences in PNNs in the insula might be driving the higher levels of aversion-resistant ethanol drinking in female mice. We did, in fact, find that female mice had more PNN deposition around neurons and increased aggrecan and brevican protein levels in the insula. Several groups have examined PNNs in different brain regions of male and female rats and mice. The number of PNNs in the cornu Ammonis-1 (CA1) region of the rat hippocampus was higher in juvenile males than females, but no significant sex differences were observed in the CA3 region or neocortex, and the sex differences in CA1 PNNs were no longer apparent in adult rats (27). Similarly, in adult rats and mice, no sex differences in PNNs in the prefrontal cortex were observed under standard housing and rearing conditions (34–36). In the basolateral amygdala (BLA), juvenile male rats had higher numbers of cells surrounded by PNNs in the left BLA and a greater percentage of PV cells with PNNs in the right BLA than females (26). The intensity of PNN staining was also higher in the medial amygdala and medial tuberal nucleus of adult male compared to female mice (25). These results are the opposite of what we observed in the insula, in which females had increased PNN staining. This discrepancy could be due to regional differences. To add another layer of complexity, PNNs are regulated by hormones in female rats and mice (37, 38).

Sex differences in the intensity of PNN staining might also be affected by the timing of brain tissue collection. We examined PNNs during the dark phase of the diurnal cycle, whereas the aforementioned studies measured PNNs during the light phase. The intensity of PNN staining by WFA has been reported to fluctuate throughout the diurnal cycle (28, 29). PNN staining was higher in the medial prefrontal cortex of rats (28) and the prefrontal cortex, hippocampus, amygdala, and other brain regions in mice (29) during the dark phase, when rodents are more active. We have not directly compared PNNs between light and dark phases but we have observed that there is no longer a sex difference in PNNs in the insula when mice are euthanized during the light phase of the cycle (data not shown). However, it is important to note that our alcohol drinking experiments are done in the dark so we collected tissue for PNNs analysis in the dark phase.

To our surprise, disrupting PNNs in the insula of female mice had no effect on aversion-resistant ethanol consumption. There are several possible explanations for this. First, this behavior may be differentially encoded anatomically in female brain. We injected ChABC into the anterior insula, but other insular subregions are involved in processing of negative information (39). The posterior insula plays a role in processing aversive states (40) and sex differences in posterior insula cortex connectivity with the bed nucleus of the stria terminalis have been observed in mice (41) and humans (42). In addition, sexually dimorphic effects of ethanol on the posterior insula to BNST circuit have also been described (41). The prefrontal cortex (PFC) is another possibility, since a PFC to nucleus accumbens circuit promotes aversion-resistant ethanol intake in rats (14). Finally, other brain regions involved in aversion should be considered, such as the habenula and rostral medial tegmental area (43).

The other explanation for the lack of effect of intra-insular ChABC in female mice on aversion-resistant ethanol consumption could be due to hormonal influences on PV interneurons. Estrogen (17β-estradiol, E2) increases the excitability of cortical PV interneurons in rats through actions at estrogen receptor β in PV neurons (44). In the rat agranular insula, approximately 32% of all PV+ neurons express estrogen receptor β (45). In addition, in the mouse hippocampus, PV interneurons have been shown to express aromatase, the enzyme that converts testosterone to E2, and blocking aromatase activity in females increased the intensity of PNNs surrounding PV interneurons (38). Interestingly, ChABC injection into the hippocampus of female mice did not affect inhibitory neurotransmission onto CA1 pyramidal cells, but was able to block the effect of an aromatase inhibitor on inhibitory currents, suggesting that when local E2 synthesis is intact, disruption of PNNs does not alter inhibitory activity in the female hippocampus (38). We hypothesize that estrogen in female insula could mask the effects of PNN removal on PV interneuron excitability and thus behavioral output. The increase in PNN staining intensity in female mouse insula might be a consequence of higher estrogen levels than in males.

These findings may also be consistent with our c-fos results. We found that induction of c-fos expression in the insula was reduced in females compared to males during quinine-adulterated ethanol consumption, and we previously demonstrated that c-fos is expressed in excitatory neurons in the insula during this behavior. If females have increased excitability of insular PV interneurons, one might expect reduced inhibition on pyramidal cells and reduced cfos expression during aversion-resistant ethanol consumption. More experiments are needed to determine if E2 modulation of PV interneurons in the insula is involved in PNN structure, the regulation of PV and pyramidal neuron excitability, and aversion-resistant ethanol consumption.

This study demonstrates that the mechanisms contributing to aversion-resistant ethanol consumption differ between males and females. When considering potential therapeutic targets to reduce compulsive alcohol drinking, sex differences need to be taken into account. Pharmacotherapies that reduce compulsive alcohol drinking in males may be ineffective in females. A greater understanding of the biology of sex differences in AUD will help in determine the most effective therapies for both sexes.

## Acknowledgements

This work was supported by grants from the National Institute on Alcohol Abuse and Alcoholism (U01 AA020912 and R01 AA027231 to AWL). The content is solely the responsibility of the authors and does not represent the official views of the National Institutes of Health.

## Notes

**Conflict of interest statement:** The authors declare no competing financial interests.

### Competing Interest Statement

The authors have declared no competing interest.

