## Supplementary figure 1 for "Sexually Dimorphic Role for Insular Perineuronal Nets in Aversion-Resistant Ethanol Consumption"

### Supplementary Figures

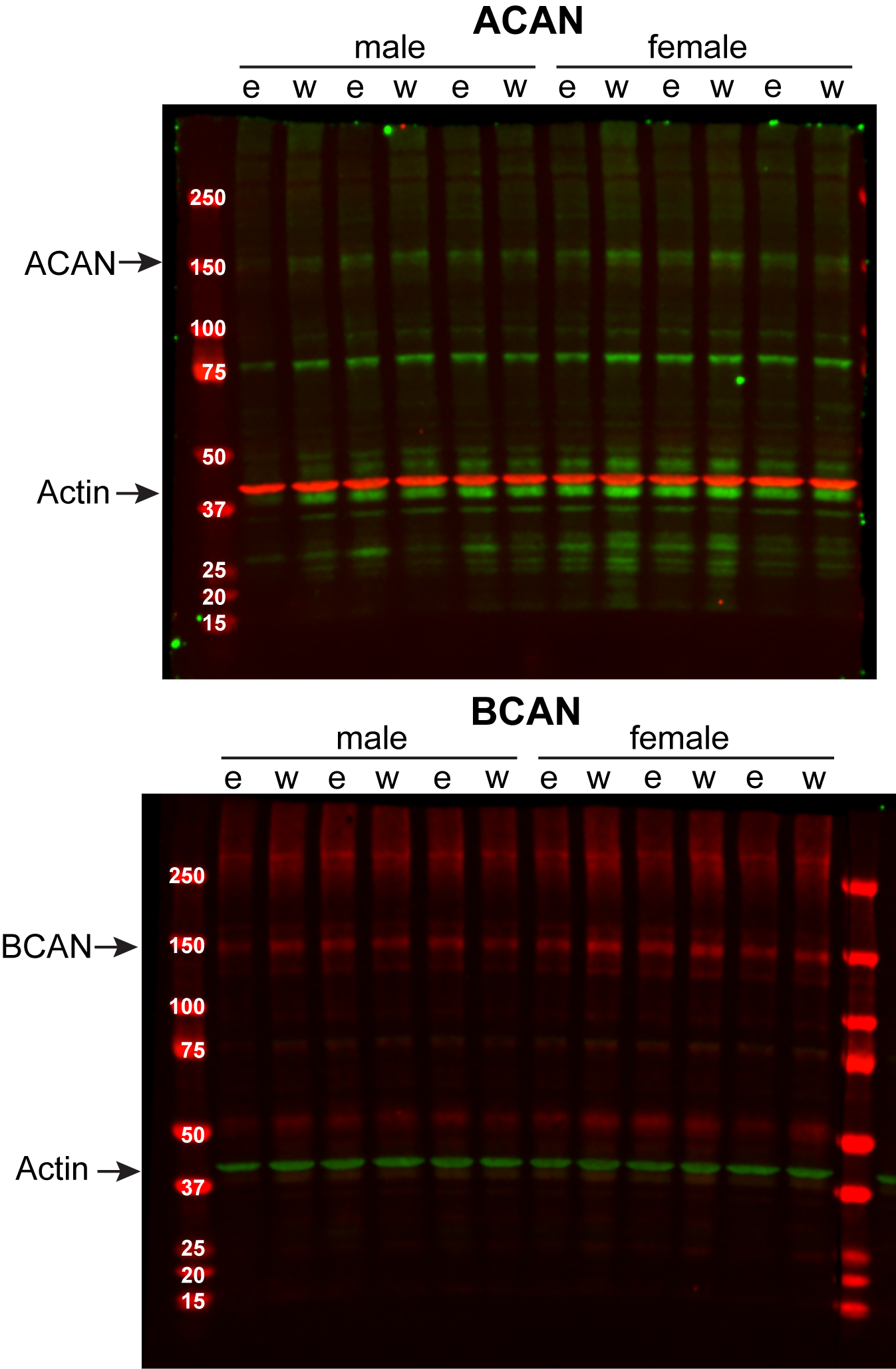

**Supplementary Figure 1.** Representative aggrecan and brevican western blot images. Aggrecan (ACAN) is the top image and brevican (BCAN) is the bottom image. Molecular weight markers were loaded in the in the first well of the gel and the molecular weight (in kDa) is indicated in white text over each band. Arrows point to ACAN, BCAN, and actin bands that were quantified. e, ethanol treated; w, water treated.
